# Stepwise C-Terminal Truncation of Cardiac Troponin T Alters Function at Low and Saturating Ca^2+^

**DOI:** 10.1101/304642

**Authors:** D. Johnson, W. Angus, J.M. Chalovich

**Author notes:** Abbreviations: EGTA, ethylene glycol-bis(β-aminoethyl ether)-N,N,N’,N’-tetraacetic acid; MOPS, 3-(N-Morpholino]-propanesulfonic acid; NEM: N-ethylmaleimide; pyrene iodoacetamide, N-(1-pyrene)iodoacetamide; Tris, tris-(hydroxymethyl)aminomethane; regulated actin, actin-tropomyosin-troponin; S1, myosin subfragment 1; TnT, troponin T; TnI, troponin I; TnC, troponin C.

## Abstract

Activation of striated muscle contraction occurs in response to Ca^2+^ binding to troponin C (TnC). The resulting reorganization of troponin repositions tropomyosin on actin and permits activation of myosin catalyzed ATP hydrolysis. It now appears that the levels of activity at both low and saturating Ca^2+^ are modulated by the C-terminal 14 amino acids of cardiac troponin T (TnT). We made a series of mutants of human cardiac troponin T, isoform 2, with deletions from the C-terminal end: Δ4, Δ6, Δ8, Δ10 and Δ14. We measured the effect of these mutations on the normalized ATPase activity at saturating Ca^2+^, the change in acrylodan tropomyosin fluorescence at low Ca^2+^, and the degree of Ca^2+^ stimulation of the rate of binding of rigor myosin S1 to pyrene-labeled actin-tropomyosin-troponin. Together, these measurements define the distribution of actin-tropomyosin-troponin among the 3 regulatory states. Results from rates of rigor S1 binding deviated from other measurements when > 8 residues of TnT were deleted. That deviation was due to increased rates of binding of rigor S1 to pyrene-labeled actin with truncated TnT at saturating Ca^2+^. Such behavior violated a key assumption in the determination of the B state by this method. Nevertheless, all methods show that as residues were removed from the C-terminus of TnT there was approximately a proportional loss of the inactive B state at low Ca^2+^ and an increase in the active M state at saturating Ca^2+^. Most of the C-terminal 14 residues of human cardiac troponin T are essential for forming the inactive B state at low Ca^2+^ and for limiting the formation of the active M state at saturating Ca^2+^.

## Introduction

Regulation of cardiac and skeletal muscle occurs through the actin binding proteins troponin and tropomyosin. At low free Ca^2+^ concentrations, these proteins reduce the ability of actin to stimulate the rate of ATP hydrolysis by myosin or the active fragments: subfragment 1 (S1) and heavy meromyosin. That inhibition appears to occur because of a decrease in the k_cat_ for ATP hydrolysis with relatively little effect on the concentration of actin-tropomyosin-troponin required for half maximum activation (1). The inhibition was not associated with a large weakening of binding of S1-ATP to actin in solution (1–3) or in rabbit psoas fibers (4) (5) although the cooperative binding of S1-ADP and rigor S1 to actin-tropomyosin-troponin are weakened at low free S1 concentrations (6). Inhibition of ATPase activity is thought to result from the presence of tropomyosin in a state that retards binding of myosin S1 to actin (7–10). That steric blocking of binding is likely restricted to “activating” forms of myosin such as rigor S1 or S1-ADP as S1-ATP and analogues of S1-ATP bind differently (11–12).

Tropomyosin is held in the inhibitory position through the concerted actions of troponin I (TnI), TnT and TnC. An increase in Ca^2+^ to permit saturation of TnC opens a hydrophobic cleft (13) to which the switch region of TnI can bind. The inhibitory region of TnI detaches from actin allowing tropomyosin to explore two other configurations. The C state (for Calcium or Closed) is the major state occupied at saturating Ca^2+^ and in the virtual absence of bound S1-ADP or rigor S1. That state, like the B state, appears to be inactive (14). In the C state, tropomyosin is partially in the actin helical groove and partially covers the high affinity myosin binding site (10). Evidence for this intermediate state came from several sources including multiple step transitions associated with the binding of myosin to actin (15–18) or dissociation of myosin from actin (19–21). Although actin in this state does not activate myosin ATPase activity, the affinity for activating types of myosin (S1-ADP and rigor S1) is increased somewhat (6) as is the rate of binding of these same species (15–16).

The other state that is populated at saturating Ca^2+^ is the M state. In this state tropomyosin is pushed more deeply into the helical groove of actin (10) because of the high affinity binding of myosin. In the M state, actin is able to stimulate the ATPase activity of myosin (21–23). The three states of actin-tropomyosin-troponin with the equilibrium constants K_B_ = [C]_eq_/[B]_eq_ and K_T_ = [M]_eq_/[C]_eq_ (16) are shown in Scheme I. Differences remain in the interpretation of these states in terms of total activity (16, 24–27) but there is agreement that the activity is proportional to the fraction of actin in the M state.

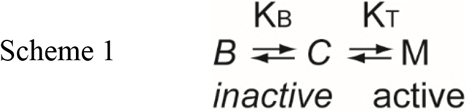

Disease causing mutations and other changes within the troponin complex alter the switching among these states (28–31). The Δ14 mutation of TnT appears to change both equilibrium constants in scheme 1. That mutant is one of two aberrant splice products resulting from a change in the splice donor sequence of exon 15 (32). Both the Δ14 and Δ28 + 7 troponin T mutations lead to an increase in activity (33–34).

Actin filaments containing Δ14 TnT have an increased rate of binding to rigor S1 at low Ca^2+^ (21), loss of the change in acrylodan-labeled tropomyosin fluorescence associated with formation of the B state (19–21) and loss of the inhibition of S1 binding at low free S1 concentrations (28). Analysis of equilibrium binding was consistent with destabilization of the inactive state (lower values of L’) without a change in binding cooperativity (28). Incorporation of Δ14 TnT into both skeletal fibers and skinned trabeculae resulted in activation at lower Ca^2+^ concentrations (28). These observations led to the conclusion that formation of the inactive B state is dependent on the last 14 residues of TnT.

The Δ14 TnT mutation also affected the behavior of actin filaments at saturating Ca^2+^. The changes observed included a doubling of the actin activated ATPase activity at saturating Ca^2+^ (21). Furthermore, troponin containing both Δ14 TnT and A8V TnC fully activated actin filaments with Ca^2+^ even in the absence of activating crossbridges. That is, the M state can be reached even in the absence of high affinity myosin binding. Indications of that possibility were shown earlier (35–36). In wild type actin filaments, the last 14 residues of TnT appear to limit the degree of activation by Ca^2+^. Neither the purpose of this limitation nor the mechanism of this limitation are understood.

The present goal is to identify key regions within the C-terminal 14 residues of human cardiac TnT that are responsible for forming the B state and for limiting the M state. The results show that the entire C-terminal region, containing 6 basic amino acid residues, is required for both effects. The final 4 residues of TnT have a slightly disproportionately large effect on activity. We also show that S1 binds more rapidly to actin in the M state than in the C state. That observation affects calculations of the fraction of B state from rates of binding of S1 to actin-tropomyosin-troponin.

## Materials and Methods

### Proteins

Isoform 2 of Human cardiac TnT in pSBETA and human cardiac TnI in pET17b were expressed and purified as described earlier (28). Human cardiac TnC in pET3d was expressed (37) and reconstituted with TnI and TnT (38). The troponin subunits were dialyzed against 1 M NaCl, 1 M urea, 5 mM MgCl_2_, 5 mM dithiothreitol, 20 mM MOPS pH 7. They were mixed in a 1:1:1.2 molar ratio and dialyzed three times, for 8h each, against the same buffer containing 6 M urea. The mixture was then dialyzed three times against the same buffer, without urea. The salt concentration was reduced by successive dialysis against the same urea-free buffer containing 0.3 M NaCl and 0.1 M NaCl. The mixture was clarified by centrifugation and applied to a Mono Q 15HR column (GE Healthcare, Pittsburgh, PA) equilibrated in the same buffer. The troponin complex was eluted with a NaCl gradient to 0.6 M. Animal use in this study was approved by the East Carolina University IACUC.

Tropomyosin was prepared from bovine cardiac left ventricles (39). Tropomyosin was labeled at cysteine 190 with acrylodan using a 10:1 molar ratio of acrylodan to tropomyosin (20). The extent of labeling (normally 70%) was determined using an extinction coefficient of 14400 M^-1^ cm^-1^ at 372 nm (40). Actin was prepared from rabbit back muscle (41) and labeled with N-(1-pyrenyl) iodoacetamide (42). Myosin was also prepared from rabbit back muscle (43) and digested with chymotrypsin to produce the soluble S1 fragment containing the enzymatic activity (44).

Tropomyosin and troponin concentrations were determined by the Lowry assay with a bovine serum albumin standard. Other protein concentrations were determined from absorbance at 280 nm with correction for light scattering at a non-absorbing wavelength. The extinction coefficients (ε^0.1%^) used for actin and S1 were 1.15 and 0.75 respectively. Molecular weights were assumed to be 42,000 (actin), 120,000 (myosin S1), 68,000 (tropomyosin), 24,000 (TnI), 35,923 (TnT) and 18,400 (TnC).

### Estimation of the M State by ATPase Rate Measurements

The initial time course of liberation of ^32^P_i_ from [γ-^32^P] ATP was measured with 3-4 time points to ensure linearity. Data were fitted to a straight line with Sigma Plot (Systat Software, Inc., San Jose, CA). Assays were run at 25 °C with 0.1 μM S1, 10 μM F-actin, and 2.2 μM tropomyosin and troponin in a buffer containing 1 mM ATP, 3 mM MgCl_2_, 34 mM NaCl, 10 mM MOPS, 1 mM dithiothreitol, 2 mM EGTA or 0.1 mM CaCl_2_, pH 7.0. Rates were normalized to the minimum, v_min_, and maximum, v_max_, values obtained under identical conditions. The normalized rates, (v_obs_-V_min_)/(V_max_-V_min_), were proportional to the fraction of actin in the M state (21). ATPase rates were corrected for the rate of hydrolysis by S1 alone.

### Relative B State Determination with Acrylodan-Labeled Tropomyosin

The fluorescence of acrylodan labeled tropomyosin is sensitive to the state of tropomyosin on actin (19, 45). The amplitude of fluorescence is proportional to the fraction of actin regulatory units in the inactive B state. The assay is illustrated in Scheme II where actin species are shown in bold letters.

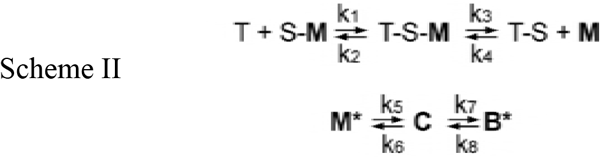

Actin filaments were stabilized in the M state by attached rigor S1 (S) under conditions of low Ca^2+^. Upon rapid binding to ATP (T), the S1 detached from M state actin with an apparent rate constant of k_3_ + k_4_ allowing actin filaments to return to the inactive C and B states with apparent rate constants k_5_ + k_6_ and k_7_ + k_8_, respectively. At low Ca^2+^ the B state predominates and the amplitude of the signal is proportional to the occupancy of the B state (19).

Actin with bound troponin, S1 and acrylodan labeled tropomyosin, were rapidly mixed with ATP at 10°C in a SF20 sequential mixing stopped-flow spectrometer (Applied Photophysics, Leatherhead, U.K.). The excitation monochrometer was set at 391 nm with a slit width of 0.5 mm. Fluorescence was measured through a long-pass filter with a cut-on midpoint of 451 nm and low and high transition points of 435 and 460 nm, respectively. The increase of the B state was followed by an increase in acrylodan-tropomyosin fluorescence.

At saturating ATP the rate of dissociation of S1 from actin was faster than the steps that followed. In most traces the detachment of S1 and the transition from the M state to the C state were too fast to be observed. The transition from the C to B state has an observed rate <10% of that for the M to C state transition. As a result, the observed transition to the B state was mono-exponential with an apparent rate constant equal to k_7_ + k_8_. Values of k_7_ + k_8_ were obtained by fitting a mono-exponential function to the time courses. The values were identical to those obtained by fitting a set of ordinary differential equations describing Scheme II to the data (19). The ratio of rate constants k_8_/k_7_ equals the ratio of state C to state B at equilibrium, that is the equilibrium constant K_B_ in Scheme I. Amplitudes were measured as the difference between the minimum and maximum fluorescence values of the mono-exponential fits.

### Estimation of B State from Kinetics of S1 binding to Pyrene-Labeled Actin

The ratio of apparent rates of binding of S1 to actin-tropomyosin-troponin at saturating Ca^2+^ to low Ca^2+^ has been used to estimate the equilibrium constant defining the equilibrium ratio of state C to state B (16):

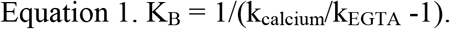

That equation requires that K_T_ << K_B_. The process is described by Scheme 3. The states of regulated actin given in bold letters. The presence of an asterisk denotes a high fluorescence state. As these measurements are normally made with rigor S1 or S1-ADP, it is assumed that the B state does not bind to regulated actin. The dashed arrows indicate that significant binding of the B state may only occur during steady-state ATP hydrolysis and not during this measurement. The values of the rate constants k_5’_and k_6’_ are only equal to k_5_ and k_6_ if the myosin S1 (S) bound equally well to actin in the C and M states. That assumption may be incorrect (see Fig. 5).

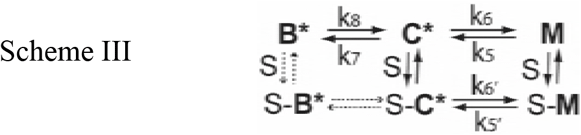

The rate of binding of S1 to pyrene labeled actin-tropomyosin-troponin was measured in the stopped flow with excitation at 365 nm with 0.5 or 1 mm slit widths and emission measured through long pass filter with a midpoint of 400 nm. Pyrene-labeled actin was stabilized with a 1:1 complex of phalloidin (Sigma-Aldrich, St. Louis, MO). The phalloidin-stabilized pyrene-labeled actin-tropomyosin-troponin complex was rapidly mixed with nucleotide-free myosin S1 in the stopped flow. Rates were measured at very low Ca^2+^ (2 mM EGTA) or at 0.5 mM Ca^2+^. Student’s t tests were performed with SigmaPlot (Systat Software, San Jose, CA) or SPSS (IBM Corp., Armonk, NY).

## RESULTS

Replacement of wild type human cardiac TnT with the Δ14 mutant reduced or eliminated the B state at low Ca^2+^ and enhanced the M state at saturating Ca^2+^ (21). Because these may represent unique regulatory functions of TnT, we sought to identify areas within the C-terminal region that are critical for these Ca^2+^ dependent effects on the distribution of actin states. We did this by measuring changes in the M and B states with a series of deletion mutants of TnT.

The effects of C-terminal TnT residues on the formation of the active M state at saturating Ca^2+^ were determined by ATPase activities. Fig.1 shows that cardiac wild type troponin-tropomyosin increased the actin stimulated rate of ATP hydrolysis by approximately 2-fold at saturating Ca^2+^. Full activation normally does not normally occur in the absence of binding of “activating” forms of myosin such as rigor S1, S1-ADP or NEM-modified S1. Fig. 1 shows that substitution of Δ4, Δ6, Δ8, Δ10 and Δ14 TnT for the wild type, increased the ATPase activity in proportion to the deletion size. A straight line between wild type and the Δ14 troponin mutations shows the expected trend if all residues contributed equally to the inhibition of M state formation.

**Figure 1.**
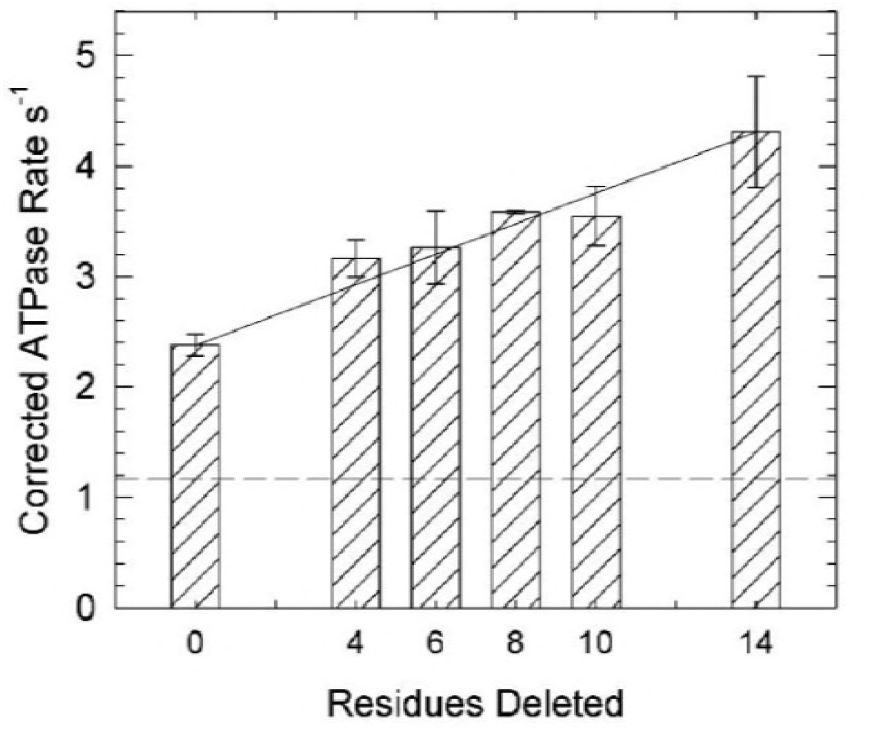
ATPase rates of myosin S1 in the presence of actin and actin-tropomyosin containing troponin with different mutants of troponin T at saturating Ca^2+^. Measurements were made at 25° C and pH 7.0 in solutions containing 1 mM ATP, 3 mM MgCl_2_, 34 mM NaCl, 10 mM MOPS, 1 mM dithiothreitol and 0.1 mM CaCl_2_. The concentrations of S1, actin, tropomyosin, and troponin were0.1, 10, 2.2, and 2.2 μM, respectively. The average rate for wild type regulated actin over all experiments was 2.4 /sec (+/-0.1). Corrected rate = average of all wild type experiments*(measured mutant rate / measured wild type rate). Error bars are standard deviation.

Slightly greater changes occurred between wild type and Δ4 and between Δ10 and Δ14. Actin filaments containing Δ14 TnT had had 1.8 times the rate of actin containing wild type troponin and 3.7 times the rate of unregulated actin (21).

Because only the M state of actin filaments stimulates myosin ATPase activity (14), the previous results can be shown in terms of fraction of M state that is formed. That is done by comparing the ATPase rates with the minimum rates and maximum possible rates at identical conditions. The minimum rate that we have observed is 0.61x the rate observed with wild type troponin at very low Ca^2+^ (30). The maximum rate was formerly determined with NEM-labeled S1 (28) or with actin filaments containing both A8V TnC and Δ14 TnT. That construct had an ATPase rate 6.5 times that of unregulated actin (21). Table I shows the population of actin in the active M state for each troponin mutant. These values are likely to be dependent on the conditions such as the ionic strength and the concentration of ATP.

**Table I.** Fraction of actin in the M state for actin filaments at saturating Ca^2+^ and containing various mutants of TnT.

|  |  |
| --- | --- |
| TnT mutant | F<sub>M</sub><sup>a</sup> |
| wild type | 0.34 |
| Δ4 | 0.45 |
| Δ6 | 0.46 |
| Δ8 | 0.50 |
| Δ10 | 0.50 |
| Δ14 | 0.62 |
| <sup>a</sup>M state = (ATPase rate) / (6.5* Unregulated actin ATPase rate) |  |

We showed earlier that acrylodan-labeled tropomyosin reports changes in the state of actin-tropomyosin-troponin allowing one to monitor the transitions from states M to C to B at low Ca^2+^ (19). We used this assay to determine the effect of deleting residues from the C-terminal of TnT on the formation of the B state at low Ca^2+^. Fig. 2 shows the time course of acrylodan tropomyosin fluorescence when S1-actin-tropomyosin-troponin was rapidly mixed with ATP. Because there was no appreciable S1 bound to actin during these reactions, the distributions observed are for free regulated actin filaments.

**Figure 2.**
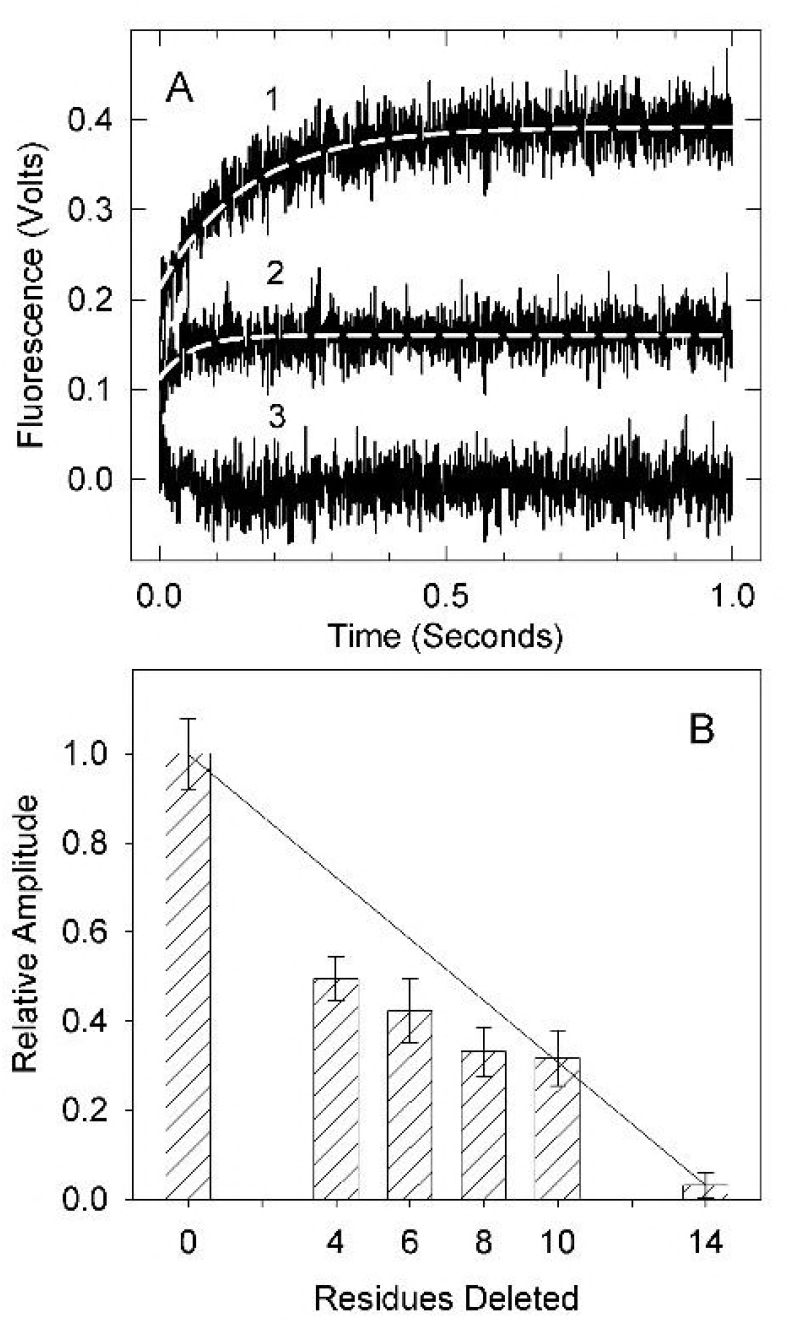
Formation of the inactive B state seen by acrylodan-tropomyosin fluorescence. A. Time courses of acrylodan fluorescence changes following the rapid detachment of myosin S1 in the absence of Ca^2+^ at 10 ^o^C. Traces shown are averages of at least three different measurements. B. Relative fluorescence amplitude for actin filaments containing wild type and truncated troponin T. The associated apparent rate constants were, 6.8, 6.9, 6.5, 7.4, 6.1 per sec for the wild type through Δ10 mutants, respectively. No rate constant could be determined for A14. Conditions: 2 μM actin, 0.86 μM tropomyosin, 1.4 μM troponin and 2 μM S1 in 20 mM MOPS, 152 mM KCl, 4 mM MgCl_2_, 1 mM dithiothreitol and 2 mM EGTA was rapidly mixed with 2 mM ATP, 20 mM MOPS, 152 mM KCl, 8 mM MgCl_2_, 1 mM dithiothreitol and 2 mM EGTA.

The decrease in fluorescence in transitioning from the M state to the C state (19) was too fast to observe in this study. Shown in Fig. 2 are the mono-exponential increases in fluorescence reflecting the slower transition from the C to the B state. Curve 1 is actin-tropomyosin with wild type troponin. The amplitude of the fluorescence increase (0.17 volts) is proportional to the amount of actin in the B state. Curve 3 shows a time course for the transition with actin regulated by troponin containing Δ14 TnT. The amplitude of the fluorescence increase in this case was near zero confirming the absence of the B state. Curve 2 is actin-tropomyosin-troponin containing the Δ10 TnT troponin truncation mutant. The Δ10 mutant had an amplitude of 0. 7 volts, intermediate to that of actin-tropomyosin containing wild type and Δ14 TnT troponin. That reduced amplitude reflects a partial loss of the B state relative to wild type.

The change in fluorescence amplitude (B state occupancy) with decreasing length of the C-terminal region of TnT is shown in Fig. 2B. Each truncation mutant had a statistically lower fluorescence amplitude than wild type regulated actin. The largest decreases in state B occupancy, occurred between WT and Δ4 and between Δ10 and Δ14. However, there is only a slight deviation from the linear curve between wild type and Δ14 TnT. Reducing the length of troponin T had no appreciable effect on the apparent rate of transition from the C state to the B state (k_7_ + k_8_).

Another way to distinguish among the states is to take advantage of their different rates of binding to myosin S1 (15, 16). In the absence of ATP, myosin S1 binds to actin-tropomyosin-troponin at a higher rate at saturating Ca^2+^ than in virtually Ca^2+^ free solution. This difference in binding occurs with either excess S1 or excess actin at pseudo first order conditions. Unlike the acrylodan-tropomyosin assay, this reports the transition of actin filaments with either some (excess actin) or total (excess S1) saturation with S1.

Fig. 3 shows time courses for binding of excess S1 to actin-tropomyosin-troponin at a very low Ca^2+^ concentration. These curves are complex because increases in the amount of S1 bound during the reaction increase both the population of the M state and the rate of binding. The presence of the B state is seen by an initial lag and a reduction in the subsequent exponential phase of binding (16, 46). This lag is thought to represent an initial blocked state population of actin filaments that do not bind to myosin (16, 47) or that bind slowly (12, 16). Binding to actin filaments containing wild type TnT, curve 1, has a lag indicative of a high population of the B state. This lag is readily seen in the inset that shows the initial part of the curve along with the exponential fit. This initial lag was largely eliminated, when wild type TnT was replaced with the Δ14 mutant (curve 2) showing that the last 14 residues of TnT are required to form the B state.

**Figure 3.**
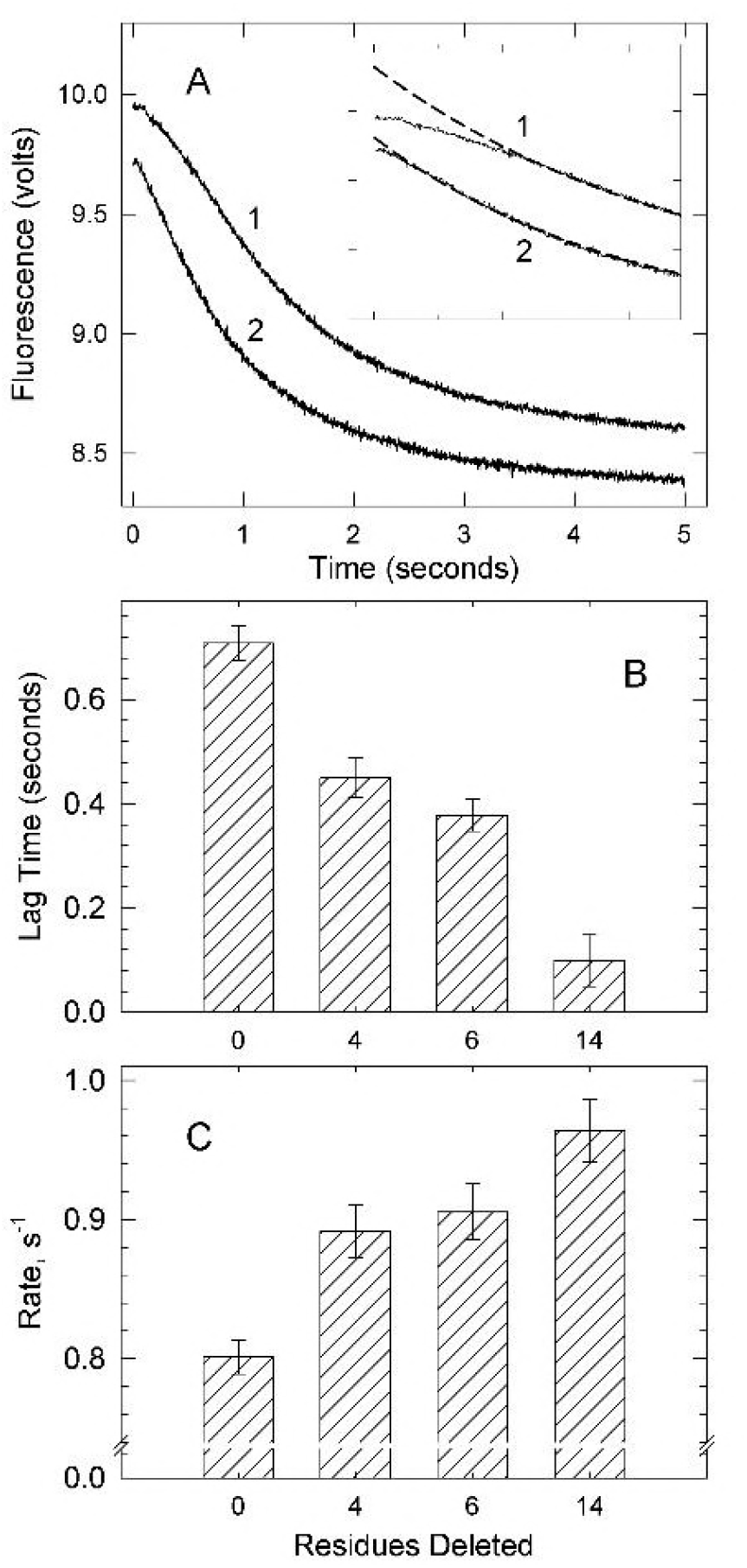
Binding of excess S1 to pyrene labeled actin filaments containing tropomyosin and troponin in the absence of ATP at very low Ca^2+^. A. Averages of > 5 time courses of S1 binding to actin filaments containing troponin with wild type troponin T, curve 1 or Δ14 troponin T, curve 2. Dashed lines are double exponential fits. B) Lag time preceding the exponential phase against troponin type. C) Apparent rate constants for the major rapid exponential phase of binding against troponin variant. Error bars shown are standard deviation. Conditions: At 25°C, 0.2 μM pyrene-actin (40% labeled), 0.086 μM tropomyosin and 0.086 μM troponin was rapidly mixed with 2 μM myosin S1 in a buffer containing 152 mM KCl, 20 mM MOPS buffer pH 7, 4 mM MgCl_2_, 1 mM dithiothreitol and 2 mM EGTA.

A histogram showing the effects of shortening the C-terminal region of TnT on the lag duration is shown in Fig. 3B. The lag time was recorded at the inflection point of each trace. TnT lacking the terminal 4, 6 or 14 residues had statistically lower lags than wild type regulated actin indicating decreases in the B state population. The extent of the lag decreased and the rate of S1 binding increased as the C-terminal region of TnT was shortened. Fig. 3C shows changes in the rate of the first and major phase of the double exponential fit to the data (the slow second phase did not vary among the mutants). Troponin mutants containing Δ4, Δ6, or Δ14 TnT each had a statistically higher rate than wild type regulated actin indicating a decrease in the B state of actin. Δ14 TnT produced the highest rate of pyrene fluorescence change corresponding to the lowest initial B state population of the troponin mutants measured. Both approaches to examine the B state (Figs 2 and 3) showed a progressive loss of the B state as the C-terminal region was truncated.

Measurements of the kinetics of S1 binding to pyrene-labeled actin are less complex when actin-tropomyosin-troponin is in large excess over S1. At that condition there is little stabilization of the C and M states due to S1 binding. Fig. 4A shows examples of binding isotherms at a very low Ca^2+^ concentration. The rates of binding increased in going from actin filaments containing wild type troponin (curve 1) to those containing either Δ6 TnT (curve 2) or Δ14 TnT (curve 3). Deletions from the C-terminal region of TnT led to an increased rate of binding indicating a lower initial B state population.

**Figure 4:**
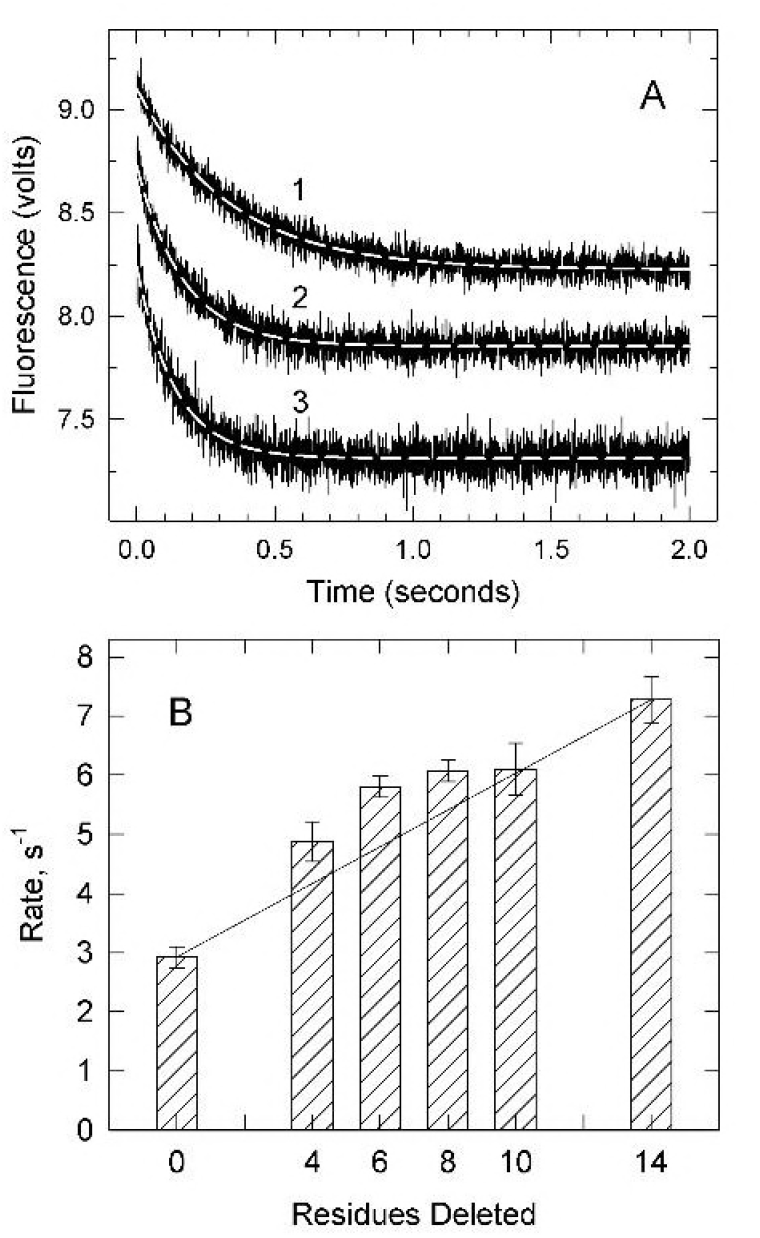
Rate of binding of S1 to an excess of pyrene labeled actin filaments containing tropomyosin and troponin in the absence of ATP at a very low Ca^2+^ concentration. A. Averaged time courses of binding to actin-tropomyosin containing wild type troponin T (curve 1); Δ6 troponin T (curve 2); and Δ14 troponin T (curve 3). Dashed lines are single exponential fits. B. Observed rate of binding of S1 to actin-tropomyosin containing troponin with different mutants of troponin T. Error bars shown are standard deviation for at least 5 experiments. Conditions: same as Fig. 3 except 4 μM pyrene-actin (40% labeled), 0.86 μM tropomyosin and 1.7 μM troponin was rapidly mixed with 0.4 μM myosin S1.

The profile of changes in rate with extent of truncation of TnT is shown in Fig. 4B. Each truncation mutant showed a statistically higher rate of fluorescence change than wild type troponin regulated actin. Deletion of the last 4 residues (GRWK) from TnT produced a large (70%) increase in the rate of binding. Additional increases were observed for subsequent deletions Δ6, Δ8 and Δ10. As in Figs. 2 and 3, the last 4 C-terminal residues appear to be particularly important.

The rate of binding of S1 to pyrene-labeled actin is thought to be at its maximum value at saturating Ca^2+^. That value is required for estimation of the B state by Equation 1. Isotherms for binding S1 to an excess of actin filaments, at saturating Ca^2+^, with wild type, Δ8 and Δ14 TnT are shown in Fig. 5A. Each curve is shown with a mono-exponential fit to the data. The rate of binding to wild type filaments was 7/sec. That value is 2.3x the wild type rate at low Ca^2+^ and is equal to the value observed for actin filaments containing Δ14 TnT at low Ca^2+^.

**Figure 5.**
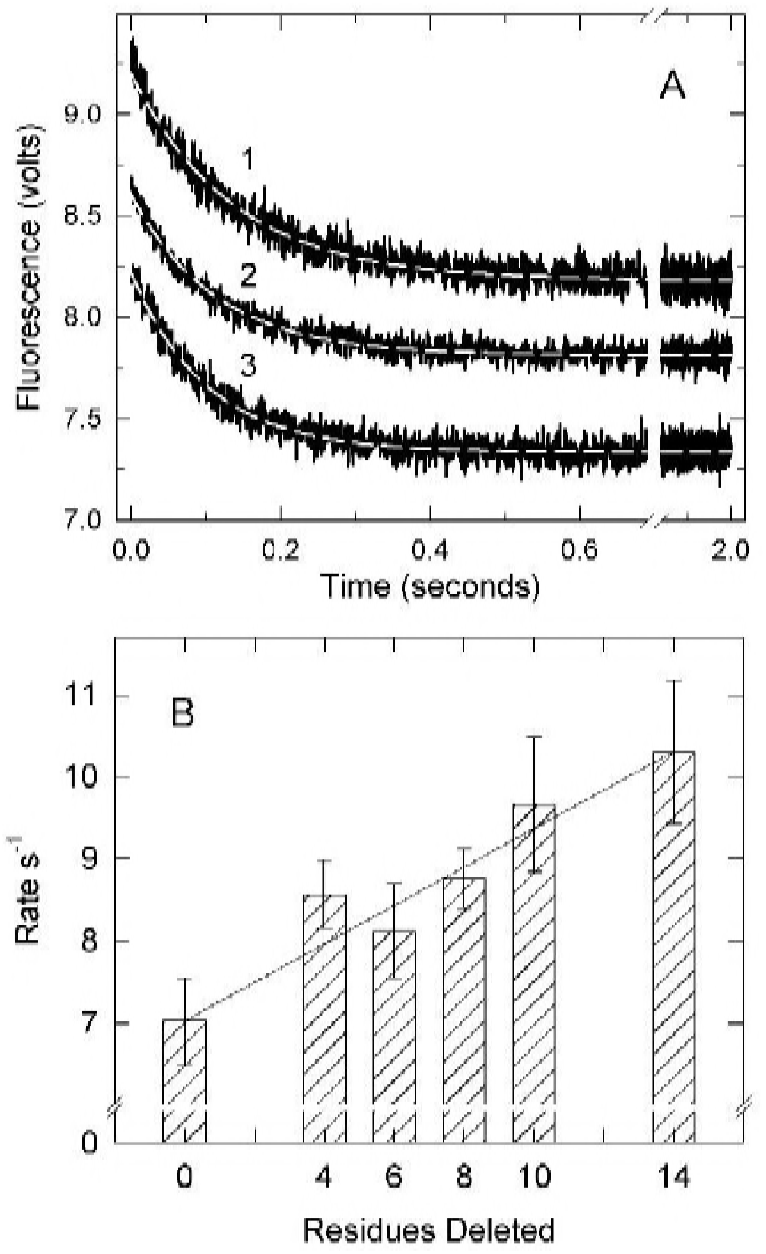
Binding of S1 to excess pyrene labeled actin filaments containing tropomyosin and troponin in the absence of ATP at saturating Ca^2+^. A. Averages of ≥5 time courses of binding to actin filaments containing troponin with wild type troponin T, curve 1; Δ8 troponin T, curve 2; and Δ14 troponin T, curve 3. Dashed lines are single exponential fits of the data. B. Observed rate of binding of S1 to actin-tropomyosin containing troponin with different mutants of troponin T. Error bars shown are standard deviation. Conditions: Same as Fig. 4 except 0.2 mM Ca^2+^ was substituted for the EGTA.

The apparent rate of binding of S1 to actin filaments at saturating Ca^2+^ increased as the C-terminal region of TnT was shortened. Fig. 5B shows that rates of binding in Ca^2+^ increased in a near linear fashion as residues were deleted from the C-terminal region of TnT. The maximum rate was with Δ14 TnT containing filaments where the apparent rate constant was about 1.4x the rate observed with wild type troponin. There was a significant increase in rate in going from wild type to Δ4 TnT with a positive deviation from the trend line.

## Discussion

The C-terminal region of TnT is essential for forming the inactive B state of actin-tropomyosin-troponin (20, 28). Furthermore, that region limits the extent of activation by Ca^2+^ (28, 31). We show here that stepwise shortening of the C-terminal region of TnT caused progressive loss of the B state at low Ca^2+^ and gain of the M state at saturating Ca^2+^. The last 4 residues (GRWK) of TnT appear to have a slightly greater effect than the remaining residues. Overall, there is nearly a linear dependence of activity on chain length.

Changes in the B state were measured by three methods. The amplitude of acrylodan tropomyosin fluorescence (Fig. 3) permits direct observation of B state levels relative to a wild type control (19, 21). The distribution of actin states obtained by that method represents the situation in the absence of bound activating crossbridges. Determining the absolute fraction of B state, by acrylodan tropomyosin fluorescence is limited by the certainty to which the wild type distribution is known. However, changes in distribution are readily measured and that is the primary goal of this study.

The other approaches for measuring the B state are based on changes in binding kinetics of rigor S1 or S1-ADP to pyrene labeled actin filaments (15–16). The absolute fraction of B state can be calculated from the ratio of rates of binding at saturating and low Ca^2+^ provided that the assumptions leading to Eq. 1 are valid: 1. the reduction in binding rates at low Ca^2+^ result from steric blocking of a fraction of potential binding sites on actin, 2. no appreciable B state exists at saturating Ca^2+^ and 3. S1 binding kinetics are the same for actin filaments in the C and M states (16). The 3^rd^ assumption was invalid in this study (Fig. 5). That is, the rate of S1 binding to pyrene actin at saturating Ca^2+^ increased as the C-terminal region of TnT was shortened giving estimated apparent rate constants of 3, 6 and 13/sec for the B, C and M states, respectively. Eq. 1 is invalid under these conditions.

The duration of the lag in binding of excess S1 to regulated filaments is a third measure of the fraction of actin filaments in the B state. The lag duration, together with acrylodan tropomyosin amplitudes and estimates from Eq. 1 are shown in Figure 6. For purposes of comparison, all data are shown as the fraction of the B state relative to that seen with wild type actin filaments. The acrylodan-tropomyosin measurements (circles) and lag durations (squares) follow a similar trend. Estimates from Eq. 1 (triangles) appeared to underestimate the fraction of B state for the mutants Δ10 through Δ14. The lag duration and acrylodan tropomyosin methods are independent of measurements in Ca^2+^ so the effects of changing distributions between the C and M state in Ca^2+^ are inconsequential. All methods showed a loss in the B state as the C-terminal region of TnT was shortened.

**Figure 6.**
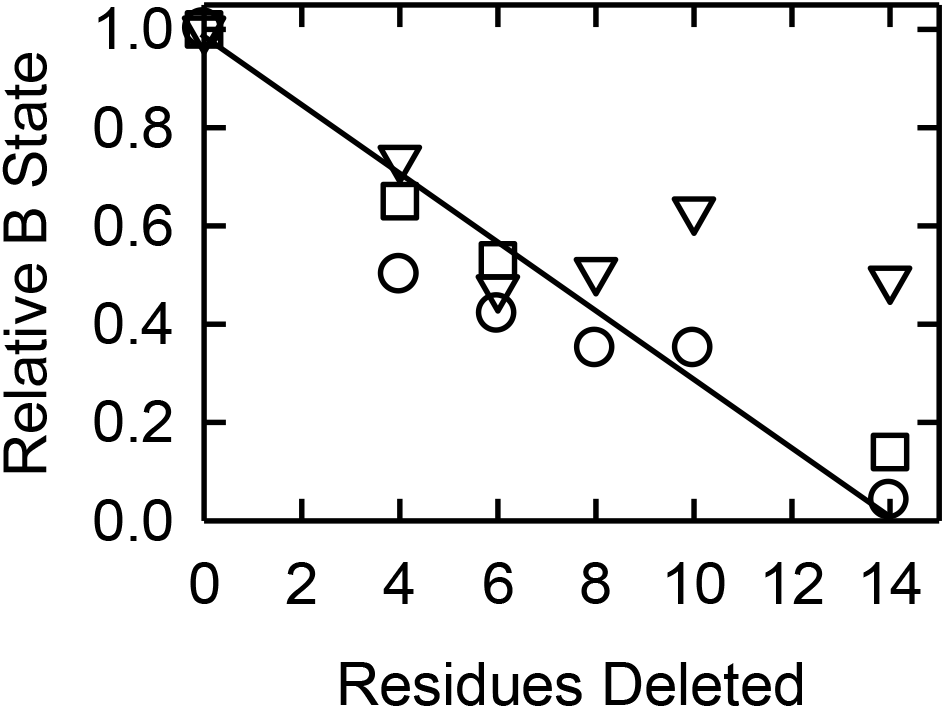
The fraction of actin filaments in the B state relative to wild type values at low Ca^2+^ for different troponin T deletion mutants. Values were calculated from acrylodan tropomyosin fluorescence amplitudes (circles), the duration of the lag in binding of excess S1 to actin (squares), and from Equation 1 (triangles). The latter were calculated using a different rate of binding in Ca^2+^ for each deletion mutant.

Although the amplitude of the B state decreased when the C-terminal region of TnT was truncated, there was no change in the apparent rate constant, k_7_ + k_8_, for transition from the C state to the B state. It is possible that the decrease in the ratio of [B]/[C] was due to both increases in k_7_ and decreases in k_8_. However, uncertainty remains because useful kinetic data could only be obtained over a limited range of [B]/[C] due to diminishing signals.

Formation of the B state at low Ca^2+^ is thought to occur because the inhibitory region (residues 137-148) and mobile domain (residues 163-210) of TnI bind to actin when the regulatory Ca^2+^ site of TnC is empty. It appears from the present results that forming this state is dependent on the entire C-terminal region of TnT. How this functions is unclear. The C-terminal region of TnT is highly basic and natively unstructured and thus invisible to X-ray and NMR analyses. Deuterium/hydrogen exchange studies showed that in the isolated troponin complex the C-terminal region of TnT exhibits rapid exchange. Ca^2+^ decreases the exchange rate of TnT at the C-terminal region of the IT helix adjacent to the C-terminal 14 residues (48). Thus, there appears to be Ca^2+^ dependent changes near the C-terminal region of TnT. The region adjacent to the C-terminus of skeletal TnT binds to TnC (49) but interactions of the last 14 residues are unknown.

The C-terminal region of TnT also retards formation of the M state at saturating Ca^2+^. The last 4 residues may have a slightly greater impact on the M state than the remainder of the C-terminal 14 residues but all residues appeared to participate in that function. At a low actin concentration, the S1 ATPase rate for actin filaments fully in the M state was 6.5 times the rate seen with S1 and pure actin (no tropomyosin or troponin) (21). The presence of troponin and tropomyosin significantly enhances actin activation at saturating Ca^2+^ (22, 28, 35, 36, 50-52). That activation beyond actin alone is most often seen with the accumulation of high affinity myosin species (rigor S1 or S1-ADP) and has been attributed to S1 pinning tropomyosin deep into the actin groove. However, troponin itself appears to modulate the position of tropomyosin on actin at saturating Ca^2+^ as well as low Ca^2+^. We show here, and in an earlier work (21), that human cardiac troponin lacking the C-terminal region of TnT gave close to 70% of full activation (i.e. 70% M state stabilization) with Ca^2+^ alone and the combination of A8V TnC and Δ14 TnT led to full activation with Ca^2+^. It may be that the large activation of ATPase activity as rigor S1 accumulates is observed only because the C-terminal region of TnT limits movement of tropomyosin into the activating position.

The importance of basic amino acid residues in the C-terminal region of TnT is reinforced by other observations. Two mutations at R278 (53–55) and two at R286 (56) result in hypertrophic cardiomyopathy. The R278C mutation leads to somewhat increased ATPase activity at both low and saturating Ca^2+^ (53) and to an increase in Ca^2+^ sensitivity for force development (53–57). That is qualitatively similar to the pattern reported here. The R278C mutation of TnT is also able to partially rescue the effects of the R145G TnI mutation at saturating Ca^2+^ (58). The R278P mutation alters the dynamics of the unstructured C-terminal region of TnT (55).

Just outside the Δ14 region is another disease causing mutation of a basic residue associated with hypertrophic and dilated cardiomyopathy. K273E TnT increased Ca^2+^ sensitivity in the in vitro motility assay (59–60) and eliminated the normal response to TnI phosphorylation (60), but did not significantly affect actomyosin ATPase activity (59).

Another deletion mutant of the C-terminal of TnT causes hypertrophic cardiomyopathy and exhibits some similar traits to the Δ14 mutation. Deletion of the 28 C-terminal residues and addition of 7 residues (Δ28+7 TnT) leads to less cooperative binding of S1 and an elevated ATPase activity at low Ca^2+^ (33). Both the Δ28 + 7 and the Δ14 mutants lowered cooperativity and increased the sensitivity of force to Ca^2+^ (34). Myocytes from transgenic mice having this mutation were also hyper-contractile (61). Yet, the position of tropomyosin at saturating and low Ca^2+^ appeared like wild type filaments (33). The affinity of troponin containing Δ28+7 TnT for actin-tropomyosin was decreased especially at low Ca^2+^ where the affinity was 22% of wild type. The authors of that study proposed that the deletion lowered the energy barrier among the states of actin so that transitions could occur more readily. In the case of Δ14 TnT the equilibrium constants appear to have changed but the rate constants between the B and C states were not greatly altered.

An involvement of both TnT and TnI in forming the B state is reasonable as there is evidence of an evolutionary and functional relationship between these subunits (58, 62). Both the C-terminal domain of TnT and the inhibitory domain of TnI (residues 137-148) emerge from the IT helix that includes residues 226-271 of human cardiac TnT (63). The inhibitory domain of TnI is highly basic, like the C-terminal region of TnT, and at 11 residues is only slightly smaller. The spatial and sequence similarity of the C-terminal region of TnT to the TnI inhibitory domain raises the possibility that the C-terminus of TnT may bind to actin and affect tropomyosin placement.

The N-terminal hypervariable region of TnT also modulates contraction (64–65). Alternative splicing at the N-terminus leads to isoforms with varied charge; those with a greater negative charge have a higher relative force in skinned fibers at low Ca^2+^ and are activated at lower Ca^2+^ concentrations (66). A higher Ca^2+^ sensitivity was also associated with a negatively charged N-terminus of TnT in chicken skeletal muscle (67). Deletion of the N-terminal region of TnT leads to a decrease in ATPase activity to 70% of wild type at low Ca^2+^ and 83% of wild type at saturating Ca^2+^ and a higher affinity for tropomyosin (68). The effects of N-terminal loss are milder than C-terminal loss and loss of charge produces decreased activity rather than increased activity as seen with C-terminal TnT truncation. While these regions have opposite charges, they do not appear to have a shared function.

The presence of an element within TnT that modulates the degree of actin activation at both low and saturating Ca^2+^ opens the possibility that interactions of the C-terminal region of TnT are regulated. It is conceivable that post translational modification of the C-terminal region of TnT, or its binding target, alters the Ca^2+^ response. Phosphorylation sites of troponin (69–70) and tropomyosin (71) are known. The C-terminal TnT residues S275 and T284 are targets for protein kinase C and the ROCKII pathway (72). The effects of phosphorylation of T284 have been studied but only in combination with phosphorylation at other sites (73–74).

## Conclusion

All of the C-terminal 14 residues of human cardiac TnT are required for forming the B state at low Ca^2+^ and for limiting the M state at saturating Ca^2+^ although the last 4 residues have a slightly greater impact than the rest. In the absence of the C-terminal residues, troponin appears to be able to stabilize tropomyosin in the active M state without much assistance from S1-ADP or rigor S1 binding to actin. This self-limitation raises the possibility that interactions of this region are regulated. Furthermore, the C-terminal region of TnT may be part of a myosin activated regulatory switch (12).

## Author Contributions

Dylan Johnson performed research, analyzed data and contributed to writing of the manuscript. William Angus designed and produced the troponin mutants. Joseph M. Chalovich designed the research, analyzed data and contributed to writing the manuscript.

## References

1. Chalovich, J. M. and E. Eisenberg. 1982. Inhibition of actomyosin ATPase activity by troponin-tropomyosin without blocking the binding of myosin to actin. The Journal of biological chemistry 257:2432–2437.

2. Chalovich, J. M., P. B. Chock, and E. Eisenberg. 1981. Mechanism of action of troponin-tropomyosin:inhibition of actomyosin ATPase activity without inhibition of myosin binding to actin. The Journal of biological chemistry 256:575–578.

3. El-Saleh, S. C. and J. D. Potter. 1985. Calcium-insensitive binding of heavy meromyosin to regulated actin at physiological ionic strength. Journal of Biological Chemistry 260:14775–14779.

4. Kraft, T., J. M. Chalovich, L. C. Yu, and B. Brenner. 1995. Parallel inhibition of active force and relaxed fiber stiffness by caldesmon fragments at physiological ionic strength and temperature conditions: Additional evidence that weak cross-bridge binding to actin is an essential intermediate for force generation. Biophys J 68:2404–2418.

5. Brenner, B., M. Schoenberg, J. M. Chalovich, L. E. Greene, and E. Eisenberg. 1982. Evidence for cross bridge attachment in relaxed muscle at low ionic strength. Proc Natl Acad Sci U S A 79:7288–7291.

6. Greene, L. E. and E. Eisenberg. 1980. Cooperative binding of myosin subfragment-1 to the actin-troponin-tropomyosin complex. Proc Natl Acad Sci U S A 77:2616–2620.

7. Huxley, H. E., R. M. Simmons, A. R. Faruqi, M. Kress, J. Bordas, and M. H. J. Koch. 1981. Millisecond time-resolved changes in x-ray reflections from contracting muscle during rapid mechanical transients, recorded using synchrotron radiation. Proceedings of the National Academy of Sciences of the United States of America 78:2297–2301.

8. Spudich, J. A., H. E. Huxley, and J. T. Finch. 1972. The regulation of skeletal muscle contraction.II.Structural studies of the interaction of the tropomyosin-troponin complex with actin. Journal of Molecular Biology 72:619–632.

9. Parry, D. A. D. and J. M. Squire. 1973. Structural role of tropomyosin in muscle regulation: analysis of the X-ray diffraction patterns from relaxed and contracting muscles. Journal of Molecular Biology 75:33–55.

10. Pirani, A., C. Xu, V. Hatch, R. Craig, L. S. Tobacman, and W. Lehman. 2005. Single particle analysis of relaxed and activated muscle thin filaments. J Mol Biol 346:761–772.

11. Chalovich, J. M., L. E. Greene, and E. Eisenberg. 1983. Crosslinked myosin subfragment 1: a stable analogue of the subfragment-1.ATP complex. Proc Natl Acad Sci U S A 80:4909–4913.

12. Resetar, A. M., J. M. Stephens, and J. M. Chalovich. 2002. Troponin-tropomyosin: an allosteric switch or a steric blocker? Biophys.J. 83:1039–1049.

13. Herzberg, O., J. Moult, and M. N. James. 1986. A model for the Ca2+-induced conformational transition of troponin C. A trigger for muscle contraction. J Biol.Chem. 261:2638–2644.

14. Mathur, M. C., T. Kobayashi, and J. M. Chalovich. 2009. Some cardiomyopathy causing troponin I mutations stabilize a functional intermediate actin state. Biophys J 96:2237–2244.

15. Trybus, K. M. and E. W. Taylor. 1980. Kinetic studies of the cooperative binding of subfragment 1 to regulated actin. Proc Natl Acad Sci U S A 77:7209–7213.

16. McKillop, D. F. A. and M. A. Geeves. 1993. Regulation of the interaction between actin and myosin subfragment 1: Evidence for three states of the thin filament. Biophys. J. 65:693–701.

17. Miki, M., T. Kobayashi, H. Kimura, A. Hagiwara, H. Hai, and Y. Maeda. 1998. Ca 2+-induced distance change between points on actin and troponin in skeletal muscle thin filaments estimated by fluorescence energy transfer spectroscopy. Journal of Biochemistry 123:324–331.

18. Kimura, C., K. Maeda, Y. Maeda, and M. Miki. 2002. Ca2+S1-Induced Movement of Troponin T on Reconstituted Skeletal Muscle Thin Filaments Observed by Fluorescence Energy Transfer Spectroscopy. Journal of Biochemistry 132:93–102.

19. Borrego-Diaz, E. and J. M. Chalovich. 2010. Kinetics of regulated actin transitions measured by probes on tropomyosin. Biophys J 98:2601–2609.

20. Franklin, A. J., T. Baxley, T. Kobayashi, and J. M. Chalovich. 2012. The C-Terminus of Troponin T Is Essential for Maintaining the Inactive State of Regulated Actin. Biophys J 102:2536–2544.

21. Baxley, T., Johnson, D., Pinto, J.R. and Chalovich, J.M. 2017. Troponin C mutations partially stabilize the active state of regulated actin and fully stabilize the active state when paired with delta 14 TnT Biochemistry.

22. Bremel, R. D., J. M. Murray, and A. Weber. 1972. Manifestations of cooperative behavior in the regulated actin filament during actin-activated ATP hydrolysis in the presence of calcium. Cold Spring Harbor Symposia on Quantitative Biology 37:267–275.

23. Williams, D. L., L. E. Greene, and E. Eisenberg. 1988. Cooperative Turning on of Myosin Subfragment-1 Adenosine-Triphosphatase Activity by the Troponin Tropomyosin Actin Complex. Biochemistry 27:6987–6993.

24. Hill, T. L., E. Eisenberg, and L. Greene. 1980. Theoretical model for the cooperative equilibrium binding of myosin subfragment 1 to the actin-troponin-tropomyosin complex. Proc Natl Acad Sci U S A 77:3186–3190.

25. Hill, T. L., E. Eisenberg, and J. M. Chalovich. 1981. Theoretical models for cooperative steady-state ATPase activity of myosin subfragment-1 on regulated actin. Biophys J 35:99–112.

26. Smith, D. A. and M. A. Geeves. 2003. Cooperative Regulation of Myosin-Actin Interactions by a Continuous Flexible Chain II: Actin-Tropomyosin-Troponin and Regulation by Calcium. Biophysical Journal 84:3168–3180.

27. Mijailovich, S. M., X. Li, J. C. del Alamo, R. H. Griffiths, V. Kecman, and M. A. Geeves. 2010. Resolution and uniqueness of estimated parameters of a model of thin filament regulation in solution. Comput Biol Chem 34:19–33.

28. Gafurov, B., S. Fredricksen, A. Cai, B. Brenner, P. B. Chase, and J. M. Chalovich. 2004. The Delta14 mutation of human cardiac troponin T enhances ATPase activity and alters the cooperative binding of S1-ADP to regulated actin. Biochemistry 43:15276–15285.

29. Mathur, M. C., P. B. Chase, and J. M. Chalovich. 2011. Several cardiomyopathy causing mutations on tropomyosin either destabilize the active state of actomyosin or alter the binding properties of tropomyosin. Biochem Biophys Res Commun 406:74–78.

30. Mathur, M. C., T. Kobayashi, and J. M. Chalovich. 2008. Negative Charges at Protein Kinase C Sites of Troponin I Stabilize the Inactive State of Actin. Biophys J 94:542–549.

31. Johnson, D., M. C. Mathur, T. Kobayashi, and J. M. Chalovich. 2016. The Cardiomyopathy Mutation, R146G Troponin I, Stabilizes the Intermediate “C” State of Regulated Actin under High-and Low-Free Ca(2+) Conditions. Biochemistry 55:4533–4540.

32. Thierfelder, L., H. Watkins, C. MacRae, R. Lamas, W. McKenna, H. P. Vosberg, J. G. Seidman, and C. E. Seidman. 1994. à-tropomyosin and cardiac troponin T mutations cause familial hypertrophic cardiomyopathy: A disease of the sarcomere. Cell 77:701–712.

33. Burhop, J., M. Rosol, R. Craig, L. S. Tobacman, and W. Lehman. 2001. Effects of a cardiomyopathy-causing troponin t mutation on thin filament function and structure. The Journal of biological chemistry 276:20788–20794.

34. Nakaura, H., S. Morimoto, F. Yanaga, M. Nakata, H. Nishi, T. Imaizumi, and I. Ohtsuki. 1999. Functional changes in troponin T by a splice donor site mutation that causes hypertrophic cardiomyopathy. American Journal of Physiology: Cell Physiology 277:C225–C232.

35. Eisenberg, E. and R. R. Weihing. 1970. Effect of skeletal muscle native tropomyosin on the interaction of amoeba actin with heavy meromyosin. Nature 228:1092–1093.

36. Chalovich, J. M. and D. Johnson. 2016. Commentary: Effect of Skeletal Muscle Native Tropomyosin on the Interaction of Amoeba Actin with Heavy Meromyosin. Front Physiol 7:377.

37. Pinto, J. R., M. S. Parvatiyar, M. A. Jones, J. Liang, M. J. Ackerman, and J. D. Potter. 2009. A functional and structural study of troponin C mutations related to hypertrophic cardiomyopathy. The Journal of biological chemistry 284:19090–19100.

38. Kobayashi, T. and R. J. Solaro. 2006. Increased Ca2+ Affinity of Cardiac Thin Filaments Reconstituted with Cardiomyopathy-related Mutant Cardiac Troponin I. Journal of Biological Chemistry 281:13471–13477.

39. Smillie, L. B. 1982. Preparation and identification of alpha- and beta-tropomyosins. In Methods in Enzymology. D. W. Frederiksen and L. W. Cunningham, editors. Academic Press. New York. 234–241.

40. Hibbs, R. E., T. T. Talley, and P. Taylor. 2004. Acrylodan-conjugated Cysteine Side Chains Reveal Conformational State and Ligand Site Locations of the Acetylcholine-binding Protein. Journal of Biological Chemistry 279:28483–28491.

41. Spudich, J. A. and S. Watt. 1971. The regulation of rabbit skeletal muscle contraction. I. Biochemical studies of the interaction of the tropomyosin-troponin complex with actin and the proteolytic fragments of myosin. Journal of Biological Chemistry 246:4866–4871.

42. Kouyama, T. and K. Mihashi. 1981. Fluorimetry study of N-(1-pyrenyl)iodoacetamide-labeled F-actin: local structural change of actin protomer both on polymerization and on binding of heavy meromyosin. European Journal of Biochemistry 114:33–38.

43. Kielley, W. W. and W. F. Harrington. 1960. A model for the myosin molecule. Biochim.Biophys.Acta 41:401–421.

44. Weeds, A. G. and R. S. Taylor. 1975. Separation of subfragment-1 isozymes from rabbit skeletal muscle myosin. Nature 257:54–56.

45. Lehrer, S. S. and Y. Ishii. 1988. Fluorescence properties of acrylodan-labeled tropomyosin and tropomyosin-actin: Evidence for myosin subfragment 1 induced changes in geometry between tropomyosin and actin. Biochemistry 27:5899–5906.

46. Chen, Y., B. Yan, J. M. Chalovich, and B. Brenner. 2001. Theoretical kinetic studies of models for binding myosin subfragment-1 to regulated actin: Hill model versus Geeves model. Biophys.J. 80:2338–2349.

47. Geeves, M. A., M. Chai, and S. S. Lehrer. 2000. Inhibition of actin-myosin subfragment 1 ATPase activity by troponin I and IC: Relationship to the thin filament states of muscle. Biochemistry 39:9345–9350.

48. Kowlessur, D. and L. S. Tobacman. 2012. Significance of troponin dynamics for Ca2+-mediated regulation of contraction and inherited cardiomyopathy. The Journal of biological chemistry 287:42299–42311.

49. Blumenschein, T. M., B. P. Tripet, R. S. Hodges, and B. D. Sykes. 2001. Mapping the interacting regions between troponins T and C. Binding of TnT and TnI peptides to TnC and NMR mapping of the TnT-binding site on TnC. The Journal of biological chemistry 276:36606–36612.

50. Eisenberg, E. and W. Kielley. 1970. Native tropomyosin: effect on the interaction of actin with heavy meromyosin and subfragment-1. Biochemical and Biophysical Research Communications 40:50–56.

51. Pemrick, S. and A. Weber. 1976. Mechanism of inhibition of relaxation by N-ethylmaleimide treatment of myosin. Biochemistry 15:5193–5198.

52. Murray, J. M., M. K. Knox, C. E. Trueblood, and A. Weber. 1982. Potentiated state of the tropomyosin actin filament and nucleotide-containing myosin subfragment 1. Biochemistry 21:906–915.

53. Szczesna, D., R. Zhang, J. Zhao, M. Jones, G. Guzman, and J. D. Potter. 2000. Altered regulation of cardiac muscle contraction by troponin T mutations that cause familial hypertrophic cardiomyopathy. The Journal of biological chemistry 275:624–630.

54. Watkins, H., W. J. McKenna, L. Thierfelder, H. J. Suk, R. Anan, A. O’Donoghue, P. Spirito, A. Matsumori, C. S. Moravec, J. G. Seidman, and C. E. Seidman. 1995. Mutations in the genes for cardiac troponin T and à-tropomyosin in hypertrophic cardiomyopathy. N Engl J Med 332:1058–1064.

55. Lassalle, M. W. 2010. Defective dynamic properties of human cardiac troponin mutations. Biosci Biotechnol Biochem 74:82–91.

56. Yang, J., W. L. Liu, D. Y. Hu, T. G. Zhu, S. N. Yang, C. L. Li, L. Li, Y. H. Sun, W. L. Xie, J. G. Yang, T. C. Li, H. Bian, Q. G. Tong, and J. Xiao. 2011. [Novel mutations of cardiac troponin T in Chinese patients with hypertrophic cardiomyopathy]. Zhonghua Xin Xue Guan Bing Za Zhi 39:909–914.

57. Morimoto, S., H. Nakaura, F. Yanaga, and I. Ohtsuki. 1999. Functional consequences of a carboxyl terminal missense mutation Arg278Cys in human cardiac troponin T. Biochem Biophys Res Commun 261:79–82.

58. Brunet, N. M., P. B. Chase, G. Mihajlovic, and B. Schoffstall. 2014. Ca(2+)-regulatory function of the inhibitory peptide region of cardiac troponin I is aided by the C-terminus of cardiac troponin T: Effects of familial hypertrophic cardiomyopathy mutations cTnI R145G and cTnT R278C, alone and in combination, on filament sliding. Arch Biochem Biophys 552-553:11–20.

59. Venkatraman, G., K. Harada, A. V. Gomes, W. G. Kerrick, and J. D. Potter. 2003. Different functional properties of troponin T mutants that cause dilated cardiomyopathy. The Journal of biological chemistry 278:41670–41676.

60. Messer, A. E., C. R. Bayliss, M. El-Mezgueldi, C. S. Redwood, D. G. Ward, M. C. Leung, M. Papadaki, C. Dos Remedios, and S. B. Marston. 2016. Mutations in troponin T associated with Hypertrophic Cardiomyopathy increase Ca(2+)-sensitivity and suppress the modulation of Ca(2+)-sensitivity by troponin I phosphorylation. Arch Biochem Biophys 601:113–120.

61. Tardiff, J. C., S. M. Factor, B. D. Tompkins, T. E. Hewett, B. M. Palmer, R. L. Moore, S. Schwartz, J. Robbins, and L. A. Leinwand. 1998. A truncated cardiac troponin T molecule in transgenic mice suggests multiple cellular mechanisms for familial hypertrophic cardiomyopathy. J Clin Invest 101:2800–2811.

62. Chong, S. M. and J. P. Jin. 2009. To investigate protein evolution by detecting suppressed epitope structures. J Mol Evol 68:448–460.

63. Takeda, S., A. Yamashita, K. Maeda, and Y. Maeda. 2003. Structure of the core domain of human cardiac troponin in the Ca 2+-saturated form. Nature 424:35–41.

64. Jin, J. P. 2016. Evolution, Regulation, and Function of N-terminal Variable Region of Troponin T: Modulation of Muscle Contractility and Beyond. Int Rev Cell Mol Biol 321:1–28.

65. MacFarland, S., J. Jin, and F. Brozovich. 2002. Troponin T isoforms modulate calcium dependence of the kinetics of the cross-bridge cycle: studies using a transgenic mouse line. Arch.Biochem.Biophys 405:241.

66. Gomes, A. V., G. Guzman, J. Zhao, and J. D. Potter. 2002. Cardiac troponin T isoforms affect the Ca2+ sensitivity and inhibition of force development. Insights into the role of troponin T isoforms in the heart. The Journal of biological chemistry 277:35341–35349.

67. Ogut, O., H. Granzier, and J. P. Jin. 1999. Acidic and basic troponin T isoforms in mature fast-twitch skeletal muscle and effect on contractility. Am J Physiol 276:C1162–1170.

68. Chandra, M., D. E. Montgomery, J. J. Kim, and R. J. Solaro. 1999. The N-terminal region of troponin T is essential for the maximal activation of rat cardiac myofilaments. J Mol Cell Cardiol 31:867–880.

69. Kranias, E. G. and J. R. Solaro. 1982. Phosphorylation of troponin I and phospholamban during catecholamine stimulation of rabbit heart. Nature 298:182–184.

70. Noland, T. A., Jr. and J. F. Kuo. 1991. Protein kinase C phosphorylation of cardiac troponin I or troponin T inhibits Ca 2+-stimulated actomyosin MgATPase activity. Journal of Biological Chemistry 266:4974–4978.

71. Montgomery, K. and A. S. Mak. 1984. In vitro phosphorylation of tropomyosin by a kinase from chicken embryo. Journal of Biological Chemistry 259:5555–5560.

72. Vahebi, S., T. Kobayashi, C. M. Warren, P. P. De Tombe, and R. J. Solaro. 2005. Functional effects of Rho-kinase-dependent phosphorylation of specific sites on cardiac troponin. Circulation Research 96:740–747.

73. Sumandea, M. P., W. G. Pyle, T. Kobayashi, P. P. de Tombe, and R. J. Solaro. 2003. Identification of a Functionally Critical Protein Kinase C Phosphorylation Residue of Cardiac Troponin T. Journal of Biological Chemistry 278:35135–35144.

74. Schlecht, W., Z. Zhou, K. L. Li, D. Rieck, Y. Ouyang, and W. J. Dong. 2014. FRET study of the structural and kinetic effects of PKC phosphomimetic cardiac troponin T mutants on thin filament regulation. Arch Biochem Biophys 550-551:1–11.

